# The complete chloroplast genome of *Garcinia binucao* (Blanco) Choisy, an indigenous fruit from the Philippines

**DOI:** 10.64898/2026.03.04.709719

**Authors:** Ma. Alexandra Cacao, Judy Ann M. Muñoz, Jessa E. Coronado, Lyka Y. Aglibot, Don Emanuel M. Cardona, Lavernee S. Gueco, Jeric C. Villanueva, Cecilia Diana C. Palao, Roneil Christian S. Alonday

## Abstract

*Garcinia binucao* (Blanco) Choisy is an indigenous species endemic to the Philippines. Its fruit is traditionally used as a souring agent in local cuisine and has been reported to possess nutritional and medicinal properties. Despite its ethnobotanical significance and promising bioactive properties, the species remains underutilized. To date, no genomic resources have been published for *G. binucao*, limiting its application in food systems, genetic studies, and conservation programs. This study reports the first complete chloroplast genome of *G. binucao* from an accession conserved at the Institute of Plant Breeding, University of the Philippines Los Baños. The assembled plastome is circular with a length of 156,570 base pairs (bp). It displays the typical quadripartite structure of most angiosperms, consisting of a large single-copy (LSC) region (85,357 bp), a small single-copy (SSC) region (17,129 bp), and a pair of inverted repeats (IR), each 27,042 bp in size. A total of 128 genes were annotated, including 83 protein-coding genes, 37 transfer RNAs (tRNAs), and eight ribosomal RNAs (rRNAs), consistent with the majority of *Garcinia* species. Of the protein-coding genes, 45 are involved in photosynthesis, 28 genes for self-replication, five genes with conserved open reading frames, and five genes are associated with other functions. The GC content was 36.2%. Leucine (10.6%) was the most abundant amino acid, with a codon usage bias toward UUA. Additionally, 98 simple sequence repeats (SSRs) were detected, 88.78% consisting of A/T motifs. Phylogenomic analysis based on assembled plastome and publicly available cpDNA sequences of 17 other species in the order Malpighiales revealed that *G. indica* is the closest relative of *G. binucao*. These findings provide a framework for future research on the species, including its conservation and potential use as a genetic resource.

## Introduction

The Philippines harbors a rich diversity of fruit species, with over 400 native fruits documented across the archipelago. Of these, 173 are classified as endemic or indigenous, with notable concentrations in Regions I, IV, and X (Coronel 2011). Many of these native fruits are deeply rooted in local food traditions and ecosystems and are valued for their nutritional composition, including essential dietary fiber, protein, β-carotene, thiamine, and ascorbic acid (Department of Science and Technology-Food and Nutrition Research Institute [DOST-FNRI] 2019).

However, malnutrition remains a pressing issue. National dietary surveys show that daily fruit and vegetable consumption in the Philippines falls significantly short of the Food and Agriculture Organization-recommended 400 grams, with average intakes of just 21 grams for fruits and 58 grams for vegetables among adults (DOST-FNRI 2022). A major constraint to addressing this gap is the limited scientific understanding of indigenous fruits, particularly their genetic and genomic characteristics, which hinders their conservation, utilization, and integration into sustainable food systems.

One such underutilized species is *Garcinia binucao* (Blanco) Choisy. Commonly used in Visayan and other regional cuisines as a natural souring agent, *G. binucao* also features in traditional medicine for managing conditions such as arthritis and hypertension. Nutritional analyses have shown that its fruit pulp contains high levels of calcium, iron, magnesium, potassium, and vitamin C, while its seeds and peel are rich in crude protein, fiber, and vitamin A (Quevedo et al. 2013).

In addition to its nutritional potential, *G. binucao* has drawn scientific interest for its bioactive compounds. Phytochemical studies have identified phenolics, flavonoids, and tannins associated with antioxidant and anti-inflammatory activities (Recuenco et al. 2020). The fruit is also a natural source of hydroxycitric acid (HCA), a compound known for its anti-obesity effects, with concentrations reaching up to 4.81 g per 100 g of fruit (Bainto et al. 2018; Bagabaldo et al. 2021). Recent studies further suggest potential neuroprotective properties and pro-myogenic effects, with extracts reducing neurotoxicity and promoting myoblast growth in model organisms (Tan et al. 2024; Tantengco et al. 2018; Teresa et al. 2024).

In recent years, neglected and underutilized species have gained attention for their potential to contribute to food security, nutrition, and the Sustainable Development Goals (Talucder et al. 2024). Advances in genomics, particularly chloroplast genome (plastome) sequencing, have emerged as valuable tools in exploring plant diversity and guiding conservation strategies. The plastome’s conserved structure, maternal inheritance, and relatively slow mutation rate make it a robust marker in phylogenetics and species identification (Daniell et al. 2016).

Despite its ethnobotanical importance and promising bioactive properties, the plastome of *G. binucao* remains uncharacterized. This study reports the first complete chloroplast genome of *G. binucao* and its phylogenomic placement within the genus *Garcinia*. These genomic insights provide a foundation for further studies in conservation biology, evolutionary research, and potential future applications involving this underutilized yet valuable Philippine fruit.

## Materials and Methods

### Plant materials

The accession GB2324 was used for DNA extraction and subsequent chloroplast genome assembly. This accession is registered at the National Plant Genetic Resources Laboratory and conserved at the Old Demo Farm, Institute of Plant Breeding, University of the Philippines Los Baños (geospatial coordinates: 14.15249° N, 121.26440° E). The leaf, fruit, seed, and habit of *G. binucao* were also characterized following Beentje (2010).

### DNA Extraction

Genomic DNA was extracted from fresh young leaves of *G. binucao* using a modified CTAB-based protocol by Inglis et al. (2018). Young leaves were used due to their lower concentrations of secondary metabolites, polysaccharides, and endonucleases, which can interfere with DNA purity and yield (Lefort and Douglas 1999). These tissues are also less susceptible to oxidative browning caused by phenolic compounds (Doyle & Doyle 1990).

### Plastome Assembly, Annotation, and Visualization

High-quality genomic DNA was sent to Macrogen Inc. (South Korea) for Illumina sequencing using the TruSeq Nano DNA library preparation kit and the NovaSeq 6000 platform. Raw reads were assessed using FASTQC v1.2 (Babraham Bioinformatics & Andrews 2018). Assembly of the chloroplast genome was performed using GetOrganelle v1.7.7.1 (Jin et al. 2020), which utilized Bowtie2 v2.5.4 (Langmead et al. 2019), SPAdes v4.0.0 (Bankevich et al. 2012), and BLAST v2.16.0 for read mapping and scaffolding.

Genome circularity was confirmed with Bandage v0.9.0 (Wick et al. 2015) while annotation was performed using CPGAVAS2 (Shi et al. 2019) and GeSeq (Tillich et al. 2017). The annotated genome of was assessed using Geneious Prime v2025.1.2 to examine key features including plastome size, gene content, and GC composition. The chloroplast genome map was then visualized with OGDraw (Greiner et al. 2019).

### Genome Characterization

Simple sequence repeats (SSRs) were identified using MISA-web (Beier et al. 2017), while long repeat sequences (forward, reverse, complement, and palindromic repeats) were analyzed using REPuter (Kurtz et al. 2001) with the following parameters: maximum number of computed repeats = 50, minimum repeat size = 30, and hamming distance = 3. Codon usage bias was evaluated through Relative Synonymous Codon Usage (RSCU) analysis using MEGA v12 (Kumar et al. 2024).

### Phylogenomic Analysis

Phylogenetic relationships were inferred using the complete chloroplast genome of *G. binucao* alongside 12 publicly available sequences of *Garcinia* species. Additional representatives from Clusioids (*Calophyllum inophyllum, Calophyllum soulattri*, and *Mesua ferrea*) were included as sister groups, while *Euphorbia hypericifolia* and *Euphorbia micractina* (Euphorbiaceae) served as outgroups, given their phylogenetic position within Malpighiales (Cai et al. 2020).

All sequences were aligned using MAFFT v7.526 (Katoh & Standley 2013). Phylogenomic analysis was conducted using IQ-tree v3.0.1 (Nguyen et al. 2015). The best-fit model was determined using the built-in ModelFinder (Kalyaanamoorthy et al. 2017) of the software. The expansion and contraction of the inverted repeat (IR) regions of *G. binucao* and its closest relative with available chloroplast genomes were visualized using IRscope (Amiryousefi et al. 2018).

## Results

### Morphology of plant materials

The evergreen tree *Garcinia binucao* GB2324 has simple leaves with an entire margin and a pinnately netted venation pattern and paripinnate branching. They exhibit a caudate apex and an attenuate base. Leaf length and width measurements ranged from 14.478 to 17.018 cm and 4.826 to 7.112 cm, respectively, with petiole lengths spanning 0.127 to 1.27 cm. The adaxial surface (Figure 1a–left) is dark green, while the abaxial surface (Figure 1a–right) is a paler green.

**Figure 1.**
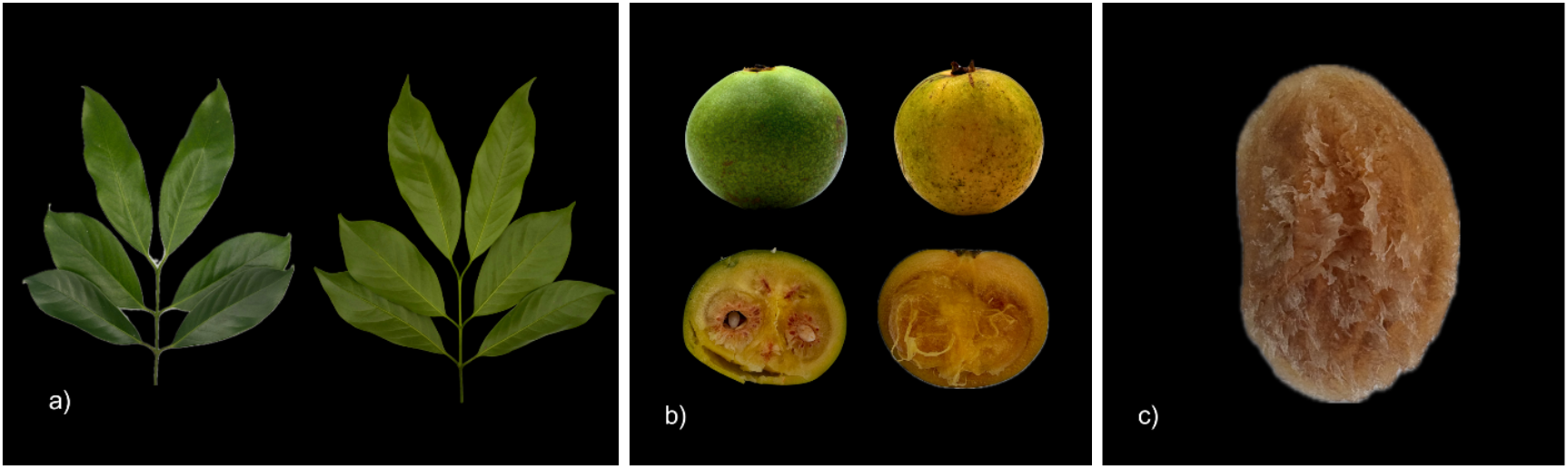
Morphological features of *Garcinia binucao* GB2324. Photographs showing a) adaxial (left) and abaxial (right) surfaces of the leaf, b) ripe and overripe fruit, c) seed, and d) whole tree (habit).

The fruit exhibits an oblate three-dimensional shape, measuring approximately 3.3 to 3.5 cm in width. Both unripe and ripe fruits appear green (Figure 1b-left), gradually transitioning to yellowish or brownish color as they over-ripen (Figure 1b-right). Each fruit typically contains 4– 6 seeds of approximately 1.5 cm in length and 1.0 cm wide, one of which is depicted in Figure 1c. The overall growth form or habit of the plant is shown in Figure 1d.

### Structure and Composition of the Plastome

The chloroplast genome of *G. binucao* is comprised of a Large Single Copy (LSC) region of 85,357 bp, a Small Single Copy (SSC) region of 17,129 bp, and two Inverted Repeat (IR) regions of 27,042 bp each, resulting in a total plastome length of 156,570 base pairs. Base composition analysis revealed 49,482 adenines (31.6%), 50,409 thymines (32.2%), 28,516 cytosines (18.2%), and 28,162 guanines (18.0%), giving a total GC content of 36.2%. The GC content varies by region: LSC (33.6%), SSC (30.2%), and IRs (42.1%). A circular representation of the plastome, including gene positions, is provided in Figure 2.

**Figure 2.**
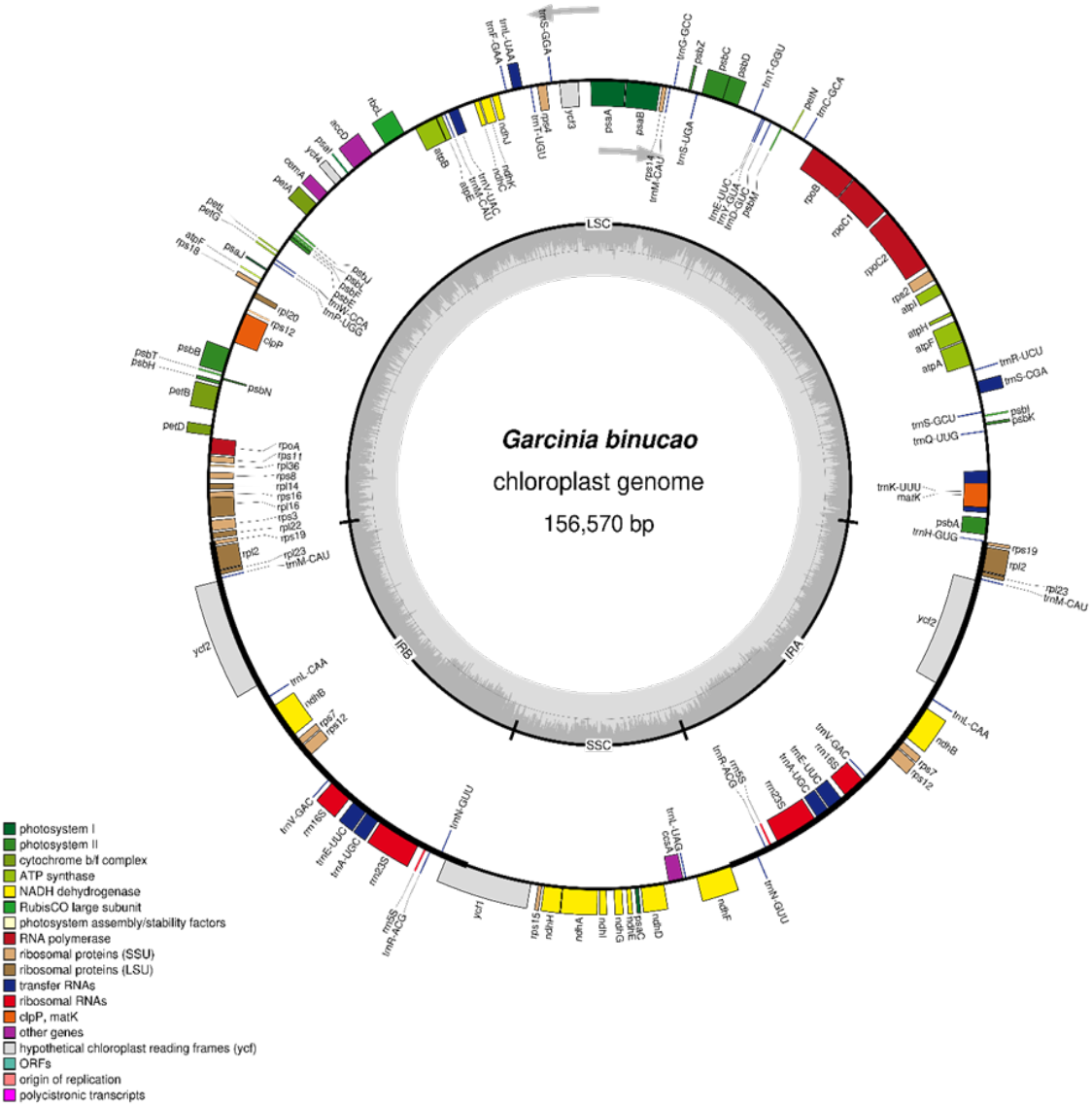
The circular chloroplast genome of *G. binucao* GB2324. As indicated by the gray arrows, genes located inside the circle are transcribed in a clockwise direction, while the genes outside the circle are transcribed counterclockwise. The GC content of the sequence is represented by the gray bars inside.

A total of 128 genes were annotated in the plastome, comprising 83 protein-coding genes (Table 1), 37 transfer RNAs (tRNAs), and eight ribosomal RNAs (rRNAs). Among the protein-coding genes, 45 are associated with photosynthesis, including six ATP synthase subunits, five photosystem I subunits, 15 photosystem II subunits, 12 NADH-dehydrogenase subunits, six cytochrome b6f complex components, and one RuBisCO large subunit.

**Table 1.**
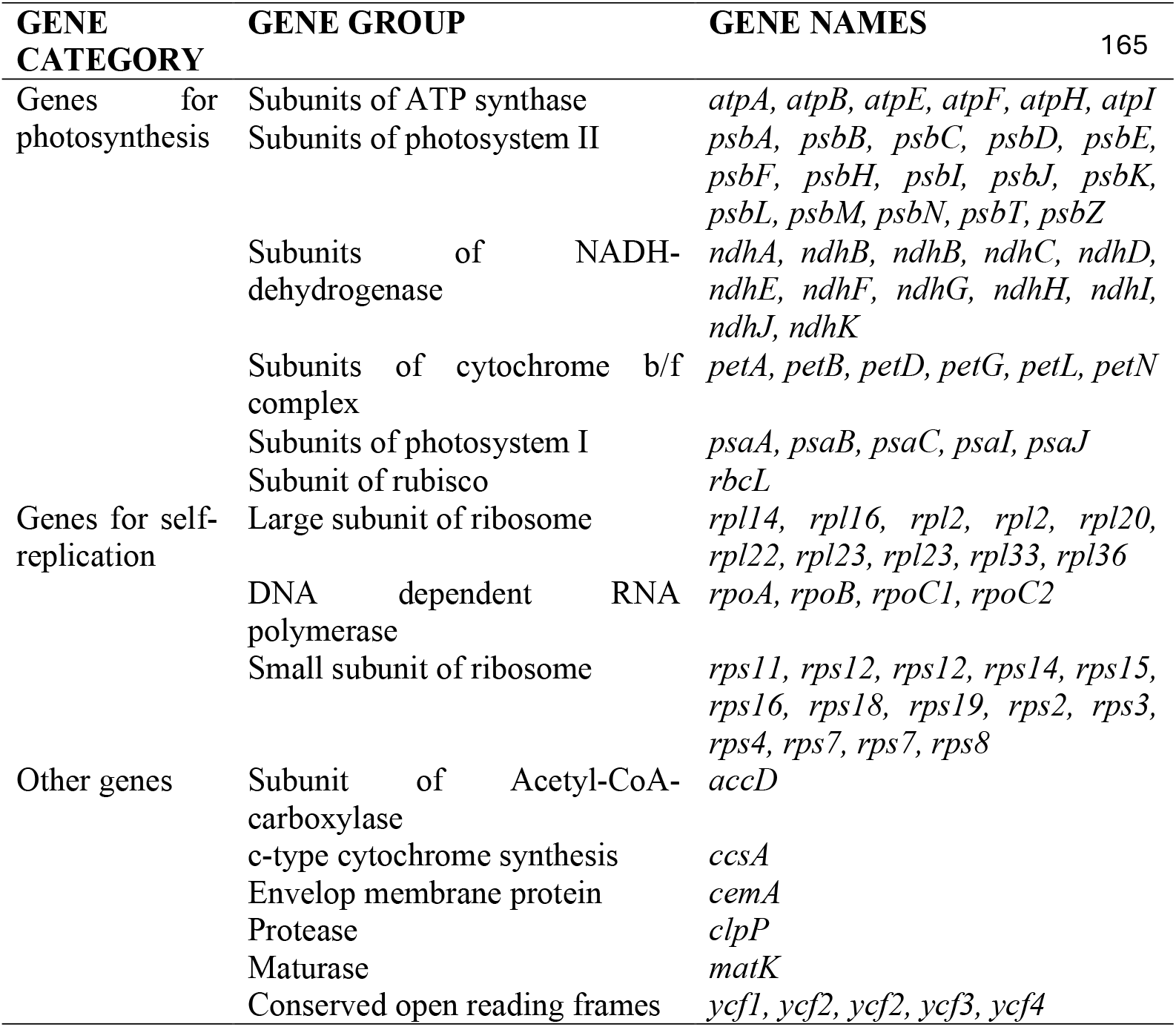
The 83 protein-coding genes of Garcinia binucao.

Additionally, 28 genes associated with genome self-replication were identified, including 10 large and 14 small ribosomal subunit genes, and four genes encoding DNA-dependent RNA polymerase subunits. Five genes with other functions were also annotated. These genes encode proteins involved in Acetyl-CoA carboxylase function, c-type cytochrome synthesis, envelope membrane integrity, protease activity, and maturase function. Furthermore, five conserved open reading frames (*ycf* genes) were identified. Gene duplication was observed for *ndhB, rpi23, ycf2, rps7*, and *rpl2*. The complete sequence and annotation of the chloroplast genome of *G. binucao* can be accessed in NCBI, accession number PX171332.

### Repetitive sequences

A total of 98 simple sequence repeats (SSRs) were identified in the chloroplast genome of *Garcinia binucao*, the majority of which were mononucleotide repeats (89), predominantly composed of A/T motifs. Dinucleotide (7) and trinucleotide (2) repeats were relatively rare, and no tetra-, penta-, or hexanucleotide motifs were detected. A/T-rich SSRs accounted for 88.78% of the total, while G/C-rich motifs comprise only 2%. These patterns are visualized in Figure 3, which highlights the dominance of A/T-rich mononucleotide SSRs. Of these repeats, 14 clusters of adjacent or overlapping units were identified.

**Figure 3.**
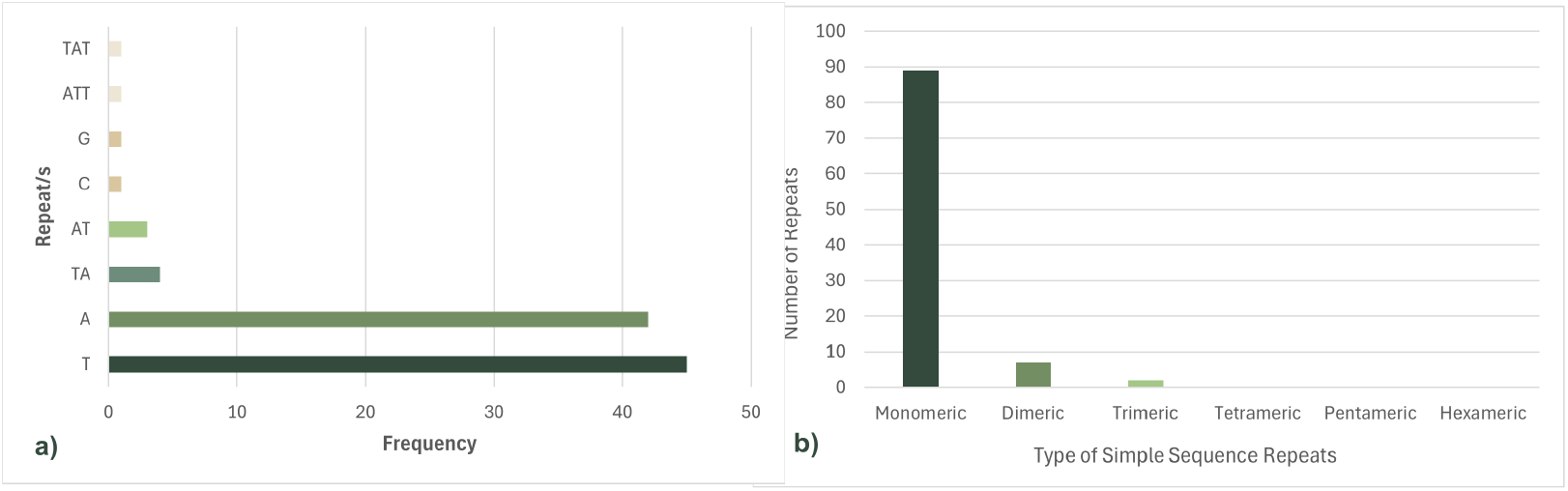
Analysis of simple sequence repeats (SSRs) in the chloroplast genome of *Garcinia binucao*. Frequency of a) specific repeat motifs, b) SSRs by the type of repeat units.

In addition to SSRs, 13 forward and 37 palindromic repeats were identified, whereas no reverse or complementary repeats were detected. The spatial distribution of SSRs and long repeats across the plastome was uneven. The LSC region contained the majority of SSRs (72.4%), followed by the IR regions (20.4%) and the SSC region (7.2%). Conversely, long repeats were most abundant in the IR regions (65.3%), followed by the LSC (18.4%) and SSC (16.3%).

### Codon usage bias

The chloroplast genome of *G. binucao* encodes a total of 26,653 codons, including stop codons. Leucine was the most frequently encoded amino acid (2,829 codons; 10.6%), followed by isoleucine (2,328; 8.7%) and serine (2,060; 7.7%). Cysteine was the least represented, with 345 codons (1.3%). A total of 214 stop codons were identified, comprising 0.8% of the codons (Supplementary Material 1).

Furthermore, Relative Synonymous Codon Usage (RSCU) analysis showed that 31 codons had values >1.0, 1 codon was equal to 1.0, and 32 codons were <1.0. The start codon AUG had an RSCU of 1.0. Among stop codons, UAA exhibited the highest bias (1.52), followed by UAG (0.88) and UGA (0.61). Notably, all the codons preferentially used for all amino acids end in A/U (Figure 4). The codon UUA (leucine) had the highest RSCU value with 1.94.

**Figure 4.**
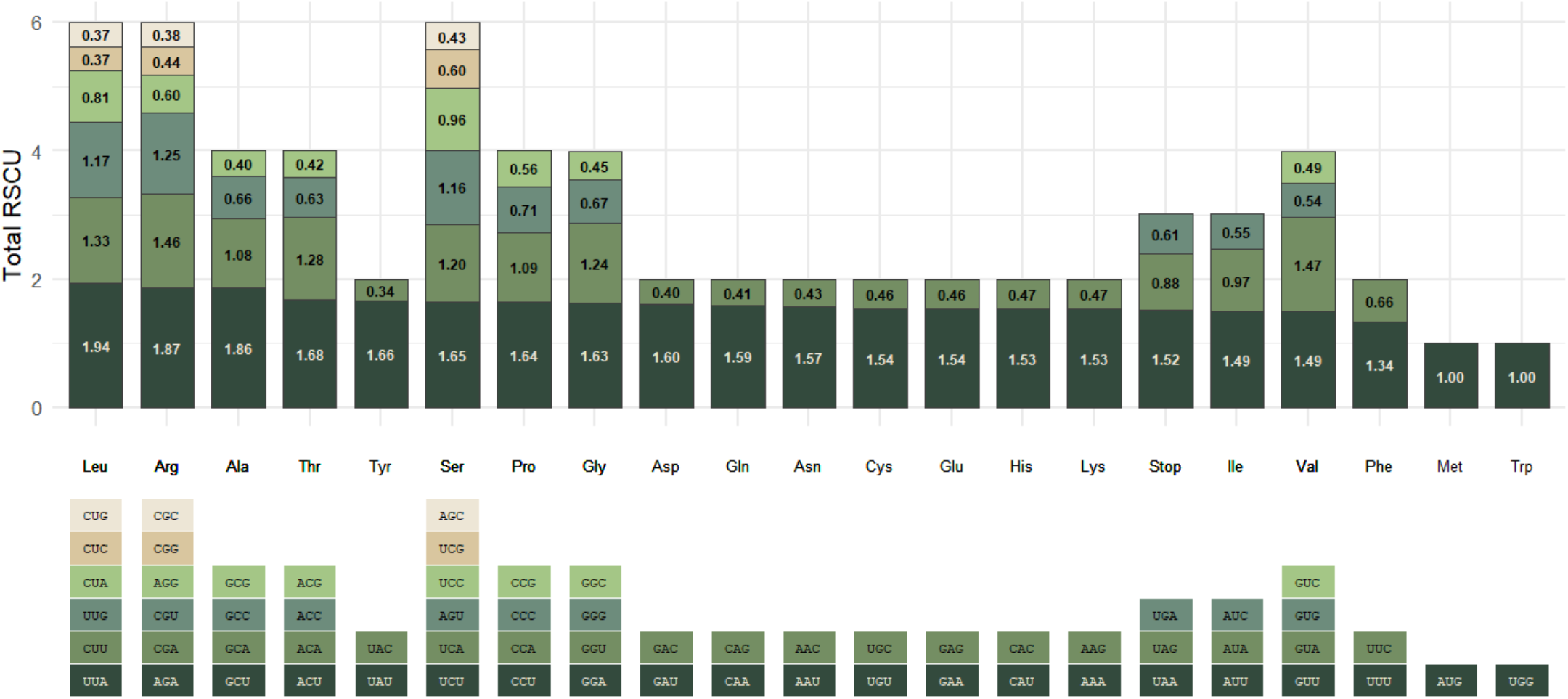
Relative synonymous codon usage (RSCU) values of codons in the chloroplast genome of *Garcinia binucao*.

### Phylogenomic analysis

Results of analysis using the assembled genome and 12 other publicly available sequences of Garcinia revealed that 1000 bootstrap replicates grouped G. binucao under the genus *Garcinia*, which has 3 clades. *G. binucao* is closest to *G. indica* and, together with *G. esculenta, G. anomala, G. gummi-gutta, G. celebica, G. esculenta, and G. oblongifolia* form one clade. The other clade consists of *G. intermedia, G. mangostana, G. pendunculata, G. xanthochymus, and G. subelliptica*, while *G. paucinervis* forms the third clade (Figure 5).

**Figure 5.**
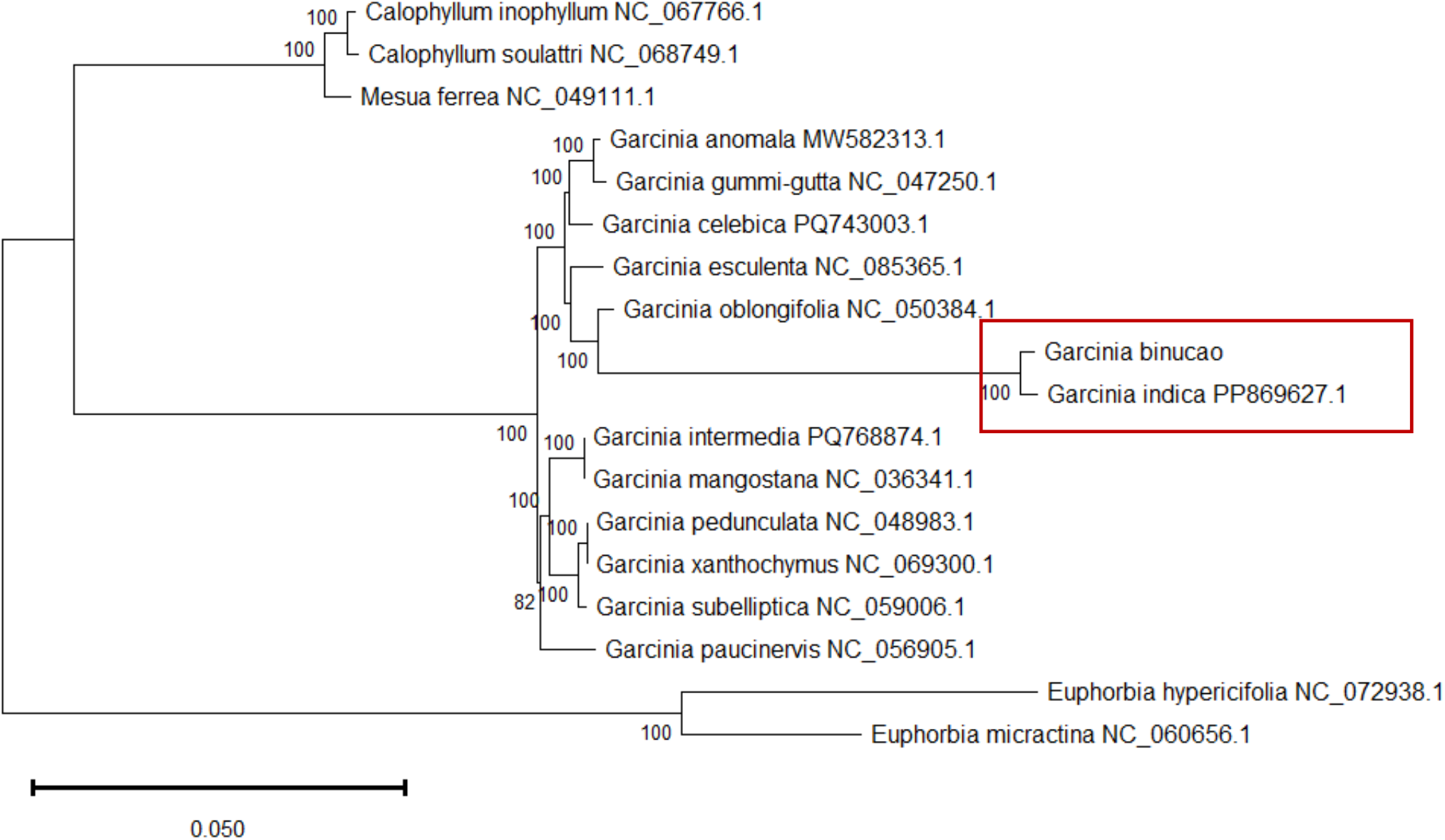
Phylogenomic tree of *Garcinia* based on complete chloroplast genomes.

### Inverted Repeat Analysis

The genes located at the four boundary regions LSC/IRb (JLB), SSC/IRb (JSB), SSC/IRa (JSA), and IRa/LSC (JLA) were *rps19, ycf1, ndhF*, and *another rps19*, respectively. At the JLB, the *rps19* gene spanned both the LSC and IRb, with 60 bp within the LSC and 219 bp in IRb. At the JSB junction, *ycf1* extended across the IRb and SSC regions, occupying 1,421 bp and 5,654 bp, respectively. The *ndhF* gene, located entirely in the SSC region, lies adjacent to the IRa boundary and measures 2,285 bp in length. Meanwhile, the other *rps19* gene was positioned at the JLA, with its sequence situated between the IRa and LSC regions. These boundary features are visualized in Figure 5 along with those of *G. indica*, showing the similarities and differences of the lengths of the genes on their boundary regions as well as the contraction and expansion, resulting in the difference in the length of the genome of the two species.

## Discussion

Most of the morphological characteristics observed in the material used for this study align with the descriptions provided by Barcelo and Barcelo (2020). The tree is evergreen, with leaves that retain their color throughout its non-fruiting period (July to February) and its fruiting season (February to June). Notably, the leaf samples used in this study were larger than those described in the reference, where leaves measuring approximately 10 cm in length and 4 cm in width were reported. In contrast, the leaves used for DNA extraction ranged from 14.478 cm to 17.018 cm in length and 4.826 to 7.112 cm in width.

The fruits from the tree where the samples were obtained were smaller (3.3 to 3.5 cm in diameter) compared to the reference fruit size (4-5 cm). Such a discrepancy may be attributed to the potential differences in the age of the tree and the timing of sample collection. Nonetheless, the characteristics of the fruits are consistent with the descriptions in the paper by Quevedo et al. (2013). Mature fruits were “green, firm, with thin skin, creamy white and less watery pulp, and a thick, woody-hard pericarp”. The endosperm was “more than half-filled and fully developed”. Upon ripening, the fruits turned “light yellow, with soft skin and pulp, retained the thick, woody-hard pericarp, and exhibited a fruity aroma,” all of which were evident in the specimens examined.

The chloroplast genome of the species exhibits the typical quadripartite structure characteristic of angiosperms, comprising the LSC, the SSC, and two IR regions. The total plastome length of 156,570 bp is consistent with the range observed among *Garcinia* species with publicly available chloroplast genome sequences. Among these, *Garcinia esculenta* possesses the shortest plastome at 155,853 bp, while *G. subelliptica* has the longest at 158,356 bp (Table 2).

**Table 2.** Comparison of the plastome size, GC content, and number of genes of Garcinia binucao and 12 other species of Garcinia. Adopted from Wang et al. (2020), Yang et al. (2019), and Yue and Shi (2021) as cited in Wee et al. (2023); Chen et al. (2022), Ma et al. (2020), Pan et al. (2025), Raju et al. (2024), Shi et al. (2024), and NCBI (2020).

| SPECIES | SIZE (BP) |  |  |  | % GC | NUMBER OF GENES |  |  |  |
| --- | --- | --- | --- | --- | --- | --- | --- | --- | --- |
|  | TOTAL | LSC | SSC | IRs |  | ALL | PCs | rRNA | tRNA |
| *G. anomala*<br>(MW582313.1) | 156,774 | 85,586 | 17,082 | 27,053 | 36.1 | 128 | 83 | 8 | 37 |
| *G. binucao*<br>(this study) | 156,570 | 85,357 | 17,129 | 27,042 | 36.2 | 128 | 83 | 8 | 37 |
| *G. esculenta*<br>(NC_085365.1) | 155,853 | 84,534 | 17,175 | 27,072 | 36.1 | 128 | 83 | 8 | 37 |
| *G. indica*<br>(PP869627.1) | 156, 891 | 85,580 | 17,181 | 27,017 | 36.2 | 126 | 86 | 8 | 32 |
| *G. gummi-gutta*<br>(NC_047250.1) | 156,202 | 84,996 | 17,088 | 27,059 | 36.1 | 127 | 83 | 8 | 36 |
| *G. mangostana*<br>(NC_036341.1) | 158,179 | 86,458 | 17,703 | 27,009 | 36.2 | 128 | 83 | 8 | 37 |
| *G. oblongifolia*<br>(NC_050384.1) | 156,577 | 85,393 | 17,064 | 27,060 | 36.2 | 128 | 83 | 8 | 37 |
| *G. paucinervis*<br>(NC_056905.1) | 157,702 | 85,989 | 17,737 | 26,988 | 36.2 | 128 | 83 | 8 | 37 |
| *G. pedunculata*<br>(NC_048983.1) | 157,688 | 85,998 | 17,656 | 27,017 | 36.2 | 128 | 83 | 8 | 37 |
| *G. subelliptica*<br>(NC_059006.1) | 158,356 | 86,220 | 17,338 | 27,399 | 36.1 | 130 | 85 | 8 | 37 |
| *G. xanthochymus*<br>(NC_069300.1) | 157,688 | 85,998 | 17,656 | 27,017 | 36.0 | 131 | 85 | 8 | 38 |

*Note:* LSC - Large Single Copy, SSC - Small Single Copy, IRs - Inverted Repeats, PC- Protein-coding, rRNA - ribosomal RNA, tRNA - transfer RNA

The plastome is predominantly composed of AT base pairs, with a GC content of 36.2%. This value falls within the expected GC range of 35-40% commonly observed in plant chloroplast genomes, such as *Broussonetia* spp. (35%; Yang et al. 2022), *Morina chinensis* (38.52%; Liu et al. 2025), and *Lirianthe* spp. (39.3%; Wu et al. 2024). Comparable GC percentages have also been reported in other *Garcinia* species (36%), suggesting a conserved base composition across the genus (Table 2). Within the structural regions of the plastome, the IRs exhibit the highest GC content, followed by the LSC region, and lastly the SSC region. This distribution is primarily attributed to the exclusive localization of the GC-rich rRNA genes within the IR regions.

The total number of annotated genes in *G. binucao* is 128, which is also consistent with other *Garcinia* species, but with minor variations: *G. indica* has the lowest number at 126, followed by *G. gummi-gutta* with 127, *G. subelliptica* with 130, and *G. xanthochymus* with the highest at 131 (Table 2). Such a difference may be attributed to gene loss or duplication events, as observed similarly in the family Solanaceae (Wang et al. 2018). Most *Garcinia* species also possess 83 protein-coding genes, although *G. indica* has 86, and both *G. subelliptica* and *G. xanthochymus* have 85. The number of tRNA genes is generally consistent at 37, with exceptions including *G. xanthochymus* (38), *G. gummi-gutta* (36), and *G. indica* (32). In contrast, the number of rRNA genes remains constant across all species, with eight genes identified for all species (Table 2).

Simple sequence repeats (SSRs) are indispensable molecular markers for phylogenetic analysis, genetic diversity, and cross-species transferability, among other molecular applications due to their high polymorphism and codominant nature (Ahmad et al. 2018). Ninety-eight SSRs were identified in the cp genome of *G. binucao*. This count is greater than that of *G. mangostana* (var. Manggis/Mesta) with 88, but is slightly lower than *G. oblongifolia*, which contains 105 SSRs (Wee et al. 2023). The predominance of A/T repeats aligns with the overall AT-rich genome composition of the species as previously discussed. TA/AT repeats account for 7.14%, while tetranucleotide motifs such as ATT/TAT represent only 2%, similar to the frequency of G/C repeats. These motifs appear only once in the genome, suggesting non-redundancy, which is advantageous for use in marker development owing to their specificity. Complex repeats were also identified in the *G. binucao* plastome, dominated by 37 palindromic repeats. This is consistent with the findings of Wee et al. (2023), who reported 25-31 palindromic repeats across *Garcinia* species. Forward repeats were the least frequent (13), which is lower than the range observed in related species (17-21). Notably, no complementary or palindromic repeats were identified, a pattern shared with *G. gummi-gutta*.

Furthermore, the RSCU value of 1.0 calculated for AUG indicates the absence of bias in its usage, as observed similarly in *G. paucinervis* (Wang et al. 2021). On the other hand, values greater than or less than 1.0 suggest that a codon is more or less frequently used in the genome, respectively. Among the stop codons, UAA was found to be 52% more frequently used than UAG and UGA, making it the preferred codon for termination. The bias in A/U as the codons’ third base was also observed in various species, such as ten species of Rutaceae in the study by Shen et al (2024). However, further analyses and comparison with other species of *Garcinia* must be conducted to fully analyze the codon usage in the plastome of the species. The high RSCU value for UUA, which encodes Leucine, suggests a strong bias towards the use of such codon, reflecting the abundance of Leucine in the plastome of the species. Data on the species’ RSCU may be used alongside other molecular data for codon optimization.

Moreover, the resulting phylogenomic tree indicates a close evolutionary relationship between *G. binucao* and *G. indica*. This relationship is strongly supported by the results of the inverted repeat boundary analysis for both species (Figure 6), which showed that identical genes occupy the junction sites of the four main regions: *rps19* at JLB, *ndhF* at JSB, *ycf1* at JSA, and *rps19* at JLA. These genes have equal lengths except for *rps19*, where a notable difference was observed. Specifically, a boundary shift at JLA marked by the contraction of the IRa and the corresponding expansion of the LSC in *G. indica* resulted in the added base pairs at the LSC from 1 bp for *G. binucao* to 80 bp for *G. indica*. The differences in the plastome lengths among species could be attributed to such events.

**Figure 6.**
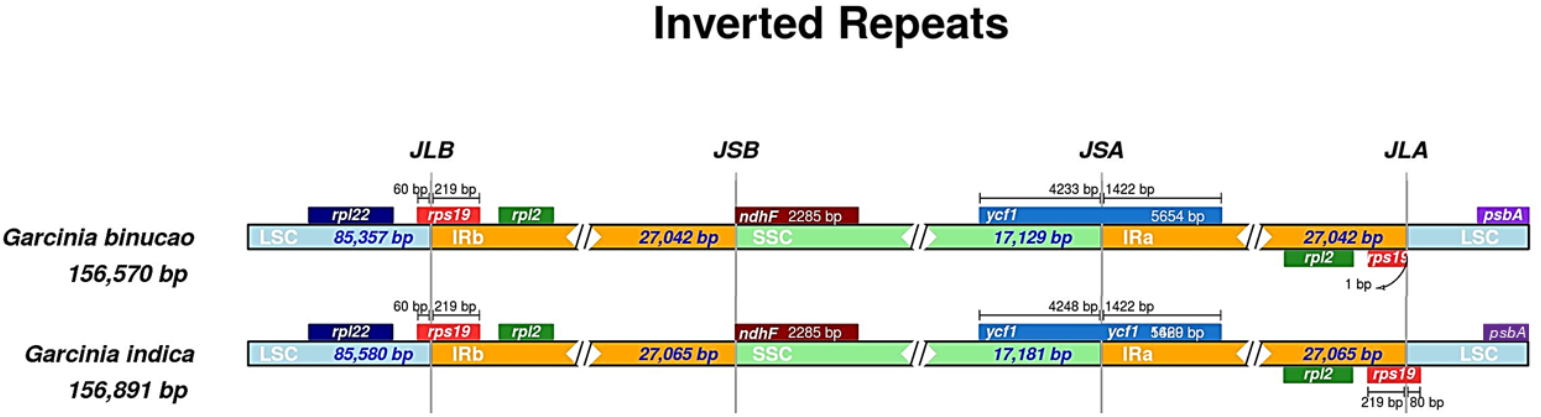
Comparison of the junction sites of Garcinia binucao and Garcinia intermedia.

The clustering of these species in a single clade alongside *G. esculenta, G. anomala, G. gummi-gutta, G. celebica*, and *G. oblongifolia* suggests a close evolutionary relationship among these species (Figure 5). This grouping could be reflective of shared environmental adaptations, morphological traits, or changes in the IR regions of these species. In contrast, a separate clade comprised of *G. intermedia, G. mangostana, G. pendunculata, G. xanthochymus*, and *G. subelliptica* indicates a distinct lineage. Lastly, *G. paucinervis* forms the third, isolated clade. The unique genomic profile of this species may be attributed to the positive selection for five key genes (*matK, psbK, rps4, rps12*, and *rps19*), which are involved in vital processes for plant development, such as protein synthesis and ribosome biogenesis (Wang et al. 2021). The grouping of the species is consistent with the phylogenetic tree of *Garcinia* spp. in the study by Shi et al. (2024)

## Conclusion

The first complete chloroplast genome of *Garcinia binucao*, an underutilized indigenous fruit species from the Philippines, was successfully assembled and annotated using leaf samples (GB2324) obtained from the Institute of Plant Breeding, University of the Philippines, Los Baños. Most of the morphological characteristics of the specimen were consistent with published descriptions of the species, supporting its taxonomic identification. The assembled plastome measured 156,570 base pairs (bp) and exhibited the typical quadripartite structure found in most angiosperms, comprising a Large Single Copy (LSC) region, a Small Single Copy (SSC) region, and two Inverted Repeats (IRs) with segment lengths comparable to those observed in other *Garcinia* species. A total of 128 genes were identified, alongside microsatellites and complex repeats, which may serve as valuable molecular markers for future genetic studies. Codon usage analysis revealed preferred codons for each amino acid, offering insights that could aid in optimizing gene expression in downstream applications. Phylogenetic analysis based on the complete chloroplast genome confirmed the placement of *G. binucao* within the genus *Garcinia*, showing a close evolutionary relationship with *G. indica*. These findings enhance our understanding of the genomic architecture and evolutionary history of *G. binucao* and may inform conservation strategies for this nutritionally and medicinally valuable indigenous species.

